# Components of alcohol use and all-cause mortality

**DOI:** 10.1101/129270

**Authors:** Sarah M. Hartz, Mary Oehlert, Amy C. Horton, Richard Grucza, Sherri L. Fisher, Karl G. Nelson, Scott W. Sumerall, P. Chad Neal, Patrice Regnier, Guoqing Chen, Alexander Williams, Jagriti Bhattarai, Bradley Evanoff, Laura J. Bierut

## Abstract

**Importance:** Current recommendations for low-risk drinking are based on drinking quantity: up to one drink daily for women and two drinks daily for men. Drinking frequency has not been independently examined for its contribution to mortality.

**Objective:** To evaluate the impact of drinking frequency on all-cause mortality after adjusting for drinks per day and binge drinking behavior.

**Design:** Two independent observational studies with self-reported alcohol use and subsequent all-cause mortality: the National Health Interview Survey (NHIS), and data from Veteran’s Health Administration clinics (VA).

**Setting:** Epidemiological sample (NHIS) and VA outpatient database (VA Corporate Data Warehouse).

**Participants:** 208,661 individuals from the NHIS interviewed between 1997 and 2009 at the age of 30 to 70 with mortality follow-up in the last quarter of 2011; 75,515 VA outpatients born between 1948 and 1968 who completed an alcohol survey in 2008 with mortality follow-up in June 2016.

**Exposures:** Quantity of alcohol use when not binging (1-2 drinks on typical day, 3-4 drinks on typical day), frequency of non-binge drinking (never, weekly or less, 2-3 times weekly, 4 or more times weekly), and frequency of binge drinking (never, less than weekly, 1-3 times weekly, 4 or more times weekly). Covariates included age, sex, race, and comorbidity.

**Main Outcomes and Measures:** All-cause mortality.

**Results:** After adjusting for binge drinking behavior, survival analysis showed an increased risk for all-cause mortality among people who typically drink 1-2 drinks four or more times weekly, relative to people who typically drink 1-2 drinks at a time weekly or less (NHIS dataset HR=1.15, 95% CI 1.06-1.26; VA dataset HR=1.31, 95% CI 1.15-1.49).

**Conclusions and Relevance:** Drinking four or more times weekly increased risk of all-cause mortality, even among those who drank only 1 or 2 drinks daily. This was seen in both a large epidemiological database and a large hospital-based database, suggesting that the results can be generalized.

## Introduction

Excessive alcohol use accounts for 9.8% of deaths among working-age adults in the United States and continues to be a leading cause of premature mortality (Stahre et al., 2014). The threshold for safe levels of alcohol use has been discussed since the mid-19^th^ century (Anstie, 1870). Currently, the National Institute of Alcohol Abuse and Alcoholism (NIAAA) and the Centers for Disease Control (CDC) recommend drinking within the U.S. Dietary guidelines, up to one drink daily for women and up to 2 drinks daily for men (National Institute on Alcohol Abuse and Alcoholism, 2016, U.S. Department of Health and Human Services and U.S. Department of Agriculture, 2015, Division of Population Health). Because of the complexity of drinking behavior itself, defining these guidelines is difficult, and therefore these recommendations have evolved over time (Stockwell and Room, 2012). Research on multiple components of drinking behavior, including number of drinking days per week, average drinks per day, and binge drinking, has informed the development of these recommendations.

There is an expansive body of literature on the relationship between alcohol use and health. Investigators have examined the impact of alcohol use on health conditions such as bodyweight, blood pressure, and stroke, as well as diseases including diabetes and multiple types of cancer (Foster and Marriott, 2006). The association between alcohol use and cardiovascular health has been studied extensively, with many studies finding evidence of a J-shaped curve (Costanzo et al., 2010b). Moderate drinkers, compared to abstainers, have a reduced risk for cardiovascular mortality, whereas heavy drinkers, compared to abstainers and moderate drinkers, have an increased risk.

A seminal study conducted by Mukamal and colleagues (2003) examined alcohol consumption and the risk of myocardial infarction among men in the Health Professionals Follow-up Study over a 12-year period. Both quantity and frequency of drinking were analyzed, along with type of alcohol consumed and changes in consumption over time. The majority of the sample reported light to moderate drinking at baseline, with only 3.5% classified as heavy drinkers. Alcohol consumption was associated with decreased risk of myocardial infarction, regardless of alcohol type, and those who reported drinking 3-4 days per week or 5-7 days per week had decreased risk compared to those who drank less than once a week, regardless of amount consumed. A moderate increase in drinking over time was also associated with decreased risk of myocardial infarction.

Several other studies investigating alcohol and cardiovascular health outcomes have also reported a protective effect for moderate drinking as measured by quantity and frequency in both men and women (e.g., Britton & Marmot, 2004; Mukamal et al., 2005; Mukamal et al. 2010; Tolstrup et al., 2006), and this protective effect has been corroborated by meta-analyses (Costanzo et al., 2010; Larson et al., 2015; Mostofsky et al., 2016; Ronksley et al., 2011).

Investigators have also examined the association between alcohol use and all-cause mortality. Some studies have replicated the J shaped curve evident with cardiovascular outcomes (Costanzo et al., 2010b, Di Castelnuovo et al., 2006, Klatsky and Udaltsova, 2007a, Ronksley et al., 2011), but other more recent studies have found no evidence for a protective effect of moderate drinking on all-cause mortality (Goulden, 2016, Stockwell et al., 2016). To further complicate matters, sex and race difference have also been reported. Sempos et al. (Sempos et al., 2003) analyzed a sample of African Americans who were followed for 19 years as part of the National Health and Nutritional Examination Survey Epidemiologic Follow-Up Study and found no protective effect for moderate drinking on all-cause mortality. Likewise, Kerr and colleagues (2011) found no protective effect of moderate drinking on all-cause mortality for African Americans when analyzing data from the National Alcohol Surveys, but the protective effect was found for whites in the same sample. With respect to sex, Klatsky and Udaltsova (2007b) reported that the protective effect of moderate drinking on all-cause mortality was stronger in women compared to men, whereas a recent meta-analysis found no protective effect at all of moderate drinking among women (Zheng et al., 2015).

The causal relationship between heavy drinking (≥ 5 drinks on a single occasion) or binge drinking (≥ 6-7 drinks on a single occasion) and detrimental health outcomes is well-established (Gonzales et al., 2014, Mokdad et al., 2004). Studies have found that heavy drinking elevates mortality risk (Plunk et al., 2014), coronary calcification (Pletcher et al., 2005), and ischemic heart disease (2007). Because an individual who drinks rarely but binges when drinking can have the same average drinks per day as an individual who drinks frequently but only drinks in moderation, it is important to characterize both average drinks per day and frequency of binging (Naimi et al., 2013, Plunk et al., 2014).

With respect to moderate drinking, however, it has been argued that protective effects on health have been overestimated and that the association, instead of being causal, is due to the methodological limitations of observational studies. The findings are so questionable, Chikritzhs et al. (2015) contend, that no protective effects should be assumed in future estimates of alcohol-related burden of disease or in national drinking guidelines. Several areas of potential bias have been identified that could lead to spurious associations, including misclassifying former drinkers as abstainers, residual or unmeasured confounding, and selection biases (Chikritzhs et al., 2009, Fillmore et al., 2007, Goulden, 2016, Klatsky and Udaltsova, 2013, Naimi et al., 2005, Naimi et al., 2017b).

Whether or not moderate alcohol consumption confers health benefits continues to be a topic of strong debate in the scientific literature (Britton and Bell, 2017, Chikritzhs et al., 2015, Mukamal et al., 2016, Naimi et al., 2017a). The current study seeks to add to the discussion regarding quantity and frequency of alcohol use and risk for all-cause mortality by analyzing two large, independent studies: an epidemiological sample from the National Health Interview Survey (NHIS) and an outpatient clinic sample from the Veteran’s Health Administration database. Specifically, we examine whether current drinkers who consume alcohol according to U.S. guidelines for safe drinking (i.e., 1-2 drinks daily) are at increased risk for all-cause mortality compared to current drinkers who drink less frequently.

## Materials and Methods

### Data

This study uses two datasets: the National Health Interview Survey (NHIS), and outpatient medical records from Veterans Health Administration clinics (VA).

#### NHIS

The NHIS is an ongoing annual survey, representative of the civilian, noninstitutionalized, household population of the United States (Minnesota Population Center and State Health Access Data Assistance Center, 2016). Data from the 1997 to 2009 administrations of the NHIS were merged to examine the relationship of drinking pattern variables with mortality data (https://ihis.ipums.org/ihis/). NHIS Data were collected via computer assisted personal interviews administered by interviewers employed and trained by the U.S. Census Bureau. Since 1997 the NHIS has included three questions to assess quantity of drinking and frequency of drinking and binge drinking in the past year, as well as two questions about lifetime alcohol use (Table 1). Respondents are told to include all types of alcoholic beverages, including liquor such as whiskey or gin, beer, wine and wine coolers. The definition of a standard drink was not provided.

**Table 1.**
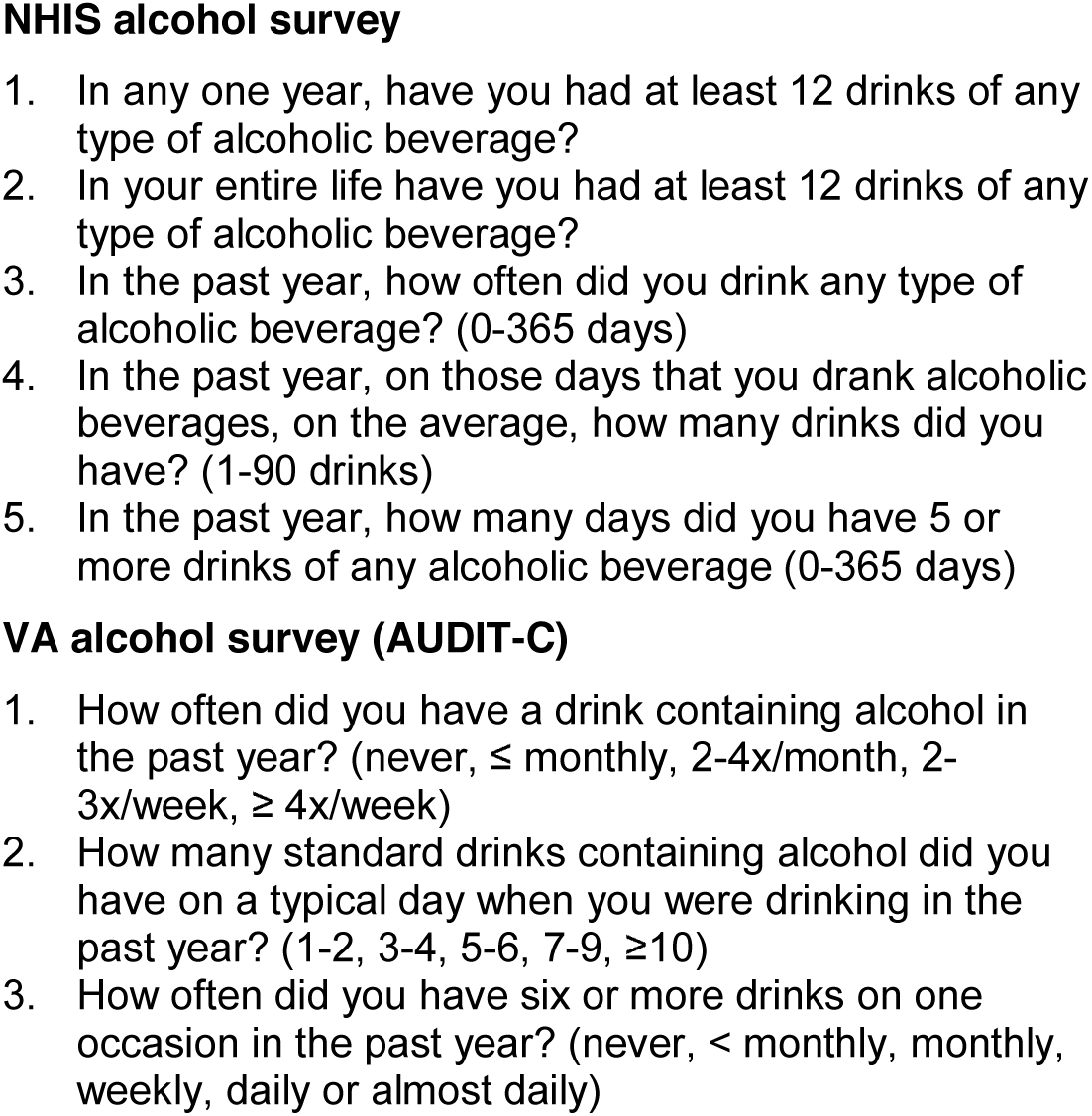
Alcohol survey

Included in these analyses were individuals who were aged 30-70 at the time of survey and had mortality follow-up in the fourth quarter of 2011. Individuals were excluded if they reported ever having cancer, a myocardial infarction, a stroke, kidney disease, or heart disease. This resulted in a total sample size of 208,661. Comorbidity was measured as self-reported medical diagnoses for select disorders coded as indicator variables (diabetes mellitus, hypertension, liver disease, peptic ulcer disease, and body mass index over 35).

#### VA Clinics

Outpatient medical records were extracted from the VA Corporate Data Warehouse (CDW) through the VA Informatics and Computing Infrastructure (VINCI). Data were extracted from the Alcohol Use Disorders Identification Test-Consumption (AUDIT-C), a brief validated screen for alcohol use disorders consisting of three items to assess quantity of drinking and frequency of drinking and binge drinking in the past year (Table 1) (Bradley et al., 2003, Bradley et al., 2007, Bush et al., 1998, Frank et al., 2008). The definition of a standard drink was not defined.

Inclusion criteria were completing an alcohol survey in 2008 (the first year the survey was introduced), date of birth between January 1, 1948 and December 31, 1968, and no past ICD-9 diagnosis of myocardial infarction, cancer, cerebrovascular disease, renal disease, or congestive heart failure. This resulted in a total sample size of 75,515. If a veteran completed more than one alcohol survey in 2008, the first administration was used for analysis. Mortality was censored at June 30, 2016. Comorbidity was measured as ICD-9 diagnoses corresponding to diabetes mellitus, peripheral vascular disease, chronic obstructive pulmonary disease, rheumatic disease, dementia, hypertension, liver disease, AIDS, and peptic ulcer disease.

#### Categories of alcohol intake

In the NHIS data, never drinkers were defined as those who had never had at least 12 drinks in their lifetime. Former drinkers were identified as those who had at least 12 drinks lifetime, but consumed no alcohol in the past year. Current drinkers were defined as those who had consumed alcohol on at least 1 day in the past year. Annual drinking frequency was converted to weekly drinking frequency by dividing by 52. For the purposes of this study, we defined binge drinking as 5 or more drinks on an occasion, which is consistent with the CDC definition of binge drinking for men.

For the VA dataset, the AUDIT-C was used to divide the sample into current non-drinkers (those who reported never having an alcoholic drink in the past year) and current drinkers (those who reported having an alcoholic drink in the past year). No information was available regarding drinking history prior to the past year. Thus, we were not able to distinguish lifetime never drinkers from former drinkers.

Among the current drinkers, patterns of alcohol use were separated into drinking quantity when not binging, frequency of non-binge drinking, and frequency of binge drinking. Drinking quantity on typical drinking days was categorized into three levels: 1-2 (moderate drinking), 3-4, and 5 or more (binge drinking). The frequency of non-binge drinking was computed by taking the overall drinking frequency (both binge drinking and non-binge drinking frequency) and subtracting the binge drinking frequency. Non-binge drinking frequency was directly computed in the NHIS data, and conservatively estimated in the VA data (see Table S1). Non-binge drinking frequency was categorized into four levels: none, weekly or less, 2-3 times weekly, and 4 or more times weekly. Binge drinking frequency was categorized into three levels: none, less than weekly, and weekly or more.

#### Data analysis

Proportional hazard analyses were completed independently in each dataset using SAS 9.4 (SAS Institute Inc., (c) 2000-2008). The outcome variable was time between the alcohol survey and either mortality or censoring of the data (fourth quarter of 2011 for NHIS data, and June 30, 2016 for VA data). In addition to the drinking pattern variables (quantity, non-binge frequency, and binge frequency), models included gender, race, age (classified as a categorical variable corresponding to 5 year bins), and comorbidity. In the NHIS data, smoking status (never, former, current) and an indicator variable for former drinker were also included in the model. A secondary analysis was run on the subset of NHIS individuals who reported never smoking. This analysis could not be run in the VA dataset because there is no systematic assessment of smoking status in the VA medical record.

In each dataset, the proportional hazards assumption was tested by evaluating the statistical significance of an interaction with time for each of the variables. To test whether there were any statistical interactions between the drinking pattern variables, an inverse stepwise regression was run with indicator variables for each three-way interaction.

## Results

The characteristics of the samples are given in Table 2. Relative to the NHIS sample, the VA sample had a shorter follow-up period with less variability, more males, more African Americans, and fewer Hispanics. In addition, the VA sample had more current drinkers, and, among those with smoking status available, there were more current smokers. As is consistent with a clinic-based sample relative to an epidemiological sample, the VA sample had a higher mortality rate, and more individuals had significant medical comorbidity.

**Table 2:** Characteristics of samples

|  | NHIS | VA |
| --- | --- | --- |
| N | 208,661 | 75,515 |
| Sample type | epidemiological | outpatient clinic |
| Year of survey | 1997-2009 | 2008 |
| Average Age at survey (SD) | 47 (11.0) | 53 (5.1) |
| Average number of years observed among non-deceased individuals (SD) | 9.1 (3.8) | 7.7 (0.22) |
| Person-years followed*** | 2,159,583 | 704,354 |
| Mortality rate (deaths per 1000 person years)*** | 7 | 13 |
| Female*** | 57% | 11% |
| Male*** | 43% | 89% |
| African American, non-Hispanic*** | 15% | 27% |
| European American, non-Hispanic*** | 65% | 52% |
| Hispanic*** | 16% | 5% |
| Other race*** | 4% | 16% |
| Current drinkers*** | 53% | 45% |
| Current non-drinkers*** | 47% | 55% |
| Never smokers | 55% | N/A |
| Current smokers | 22% | N/A |
| Former smokers | 23% | N/A |
| Presence of significant medical comorbidity*** | 42% | 54% |

NHIS: National Health Interview Survey
VA: Veteran's Health Administration clinics
SD: standard deviation
N/A: not available
\*\*\*p<0.0001 for difference between NHIS and VA

Among current drinkers, the drinking patterns between the two datasets are given in Table 3. The current drinkers in the VA sample were more likely to only binge drink (i.e., no moderate drinking) relative to the current drinkers in the NHIS sample, and to binge drink more frequently.

**Table 3:** Patterns of alcohol use across samples

|  | NHIS | VA |
| --- | --- | --- |
| Current Drinkers | 129,828 | 41,194 |
| **Average number of drinks per occasion when not bingeing** |  |  |
| 1-2 drinks | 70% | 68% |
| 3-4 drinks | 21% | 15% |
| Only binges (<u>>5 drinks</u>) | 9% | 17% |
| **Frequency of non-binge drinking** |  |  |
| Weekly or less | 62% | 57% |
| 2-3 times weekly | 19% | 16% |
| 4 or more times weekly | 11% | 10% |
| No moderate drinking (i.e. all drinking is binge drinking) | 8% | 17% |
| **Frequency of binge drinking** |  |  |
| never | 67% | 73% |
| monthly or less | 25% | 15% |
| weekly | 7% | 6% |
| 4 or more times weekly | 2% | 6% |

NHIS: National Health Interview Survey
VA: Veteran's Health Administration clinics

For the never-bingers, survival curves were plotted comparing the drinking frequency and drinking quantity strata (Figure 1; all survival curves stratified by binge drinking frequency are given in Supplementary Figure S1). Survival analysis was run in each dataset to estimate the impact of drinking quantity, non-binge drinking frequency, and binge drinking frequency on all-cause mortality. Impact of drinking patterns on mortality was estimated using hazard ratios, the multiplicative change in mortality rate due to a risk factor, adjusted for demographic characteristics and comorbidity. Hazard ratios can be conceptualized as similar to relative risk: where relative risk refers to the change in probability of a disease due to a particular risk factor, hazard ratios refer to the change in the probability of death at any given time point due to a risk factor, the assumption being that the impact of this risk factor does not vary with time (referred to as the assumption of proportional hazards). Therefore, a hazard ratio of 2 for a particular risk factor indicates that this factor doubles the risk of death at any particular time.

**Figure 1:**
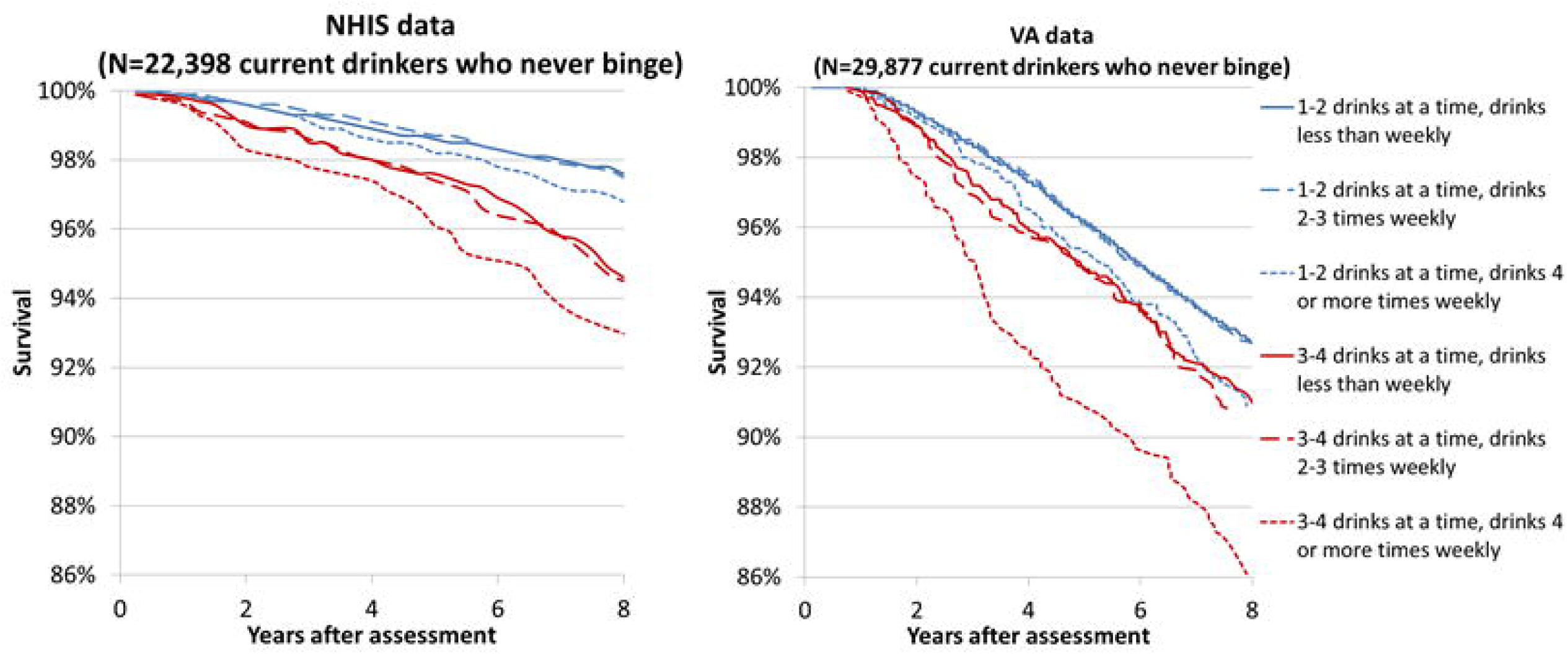
Survival curves among current drinkers who never binge.

Marginal adjusted hazard ratios for three components of drinking are given in Table 4. Because there were no statistically significant two- or three-way interactions between the drinking quantity, non-binge frequency, and binge frequency categories, each hazard ratio for an individual drinking pattern was compiled using the marginal estimates for the categories. For the never bingers, the hazard ratios for individual drinking patterns with 95% confidence intervals are graphically represented in Figure 2 (see Supplementary Figure S2 for estimated hazard ratios in all drinking strata). Among non-drinkers in the NHIS sample, former drinkers had a higher mortality rate than never drinkers (HR 1.07, p=0.03, adjusted for age, sex, race, smoking status, and comorbidity).

**Table 4:** Adjusted, marginal hazard ratios for components of drinking. Hazard ratios are from a single proportional hazards regression analysis adjusted for sex, age, race and comorbidity.

|  | NHIS |  | VA |  |
| --- | --- | --- | --- | --- |
|  | HR | 95% CI | HR | 95% CI |
| **Average number of drinks when not bingeing** |  |  |  |  |
| 1-2 drinks |  | REFERENCE |  |  |
| 3-4 drinks | 1.21 | (1.12-1.29) | 1.36 | (1.16-1.59) |
| **frequency of non-binge drinking** |  |  |  |  |
| weekly or less |  | REFERENCE |  |  |
| 2– 3x/week | 0.99 | (0.93-1.06) | 1.10 | (0.98-1.24) |
| 4 or more times weekly | 1.15 | (1.06-1.26) | 1.31 | (1.15-1.49) |
| **Frequency of binge drinking** |  |  |  |  |
| Never |  | REFERENCE |  |  |
| Monthly or less | 0.99 | (0.92-1.06) | 1.19 | (1.04-1.35) |
| 2x/month – 3x/week | 1.28 | (1.16-1.42) | 1.43 | (1.20-1.71) |
| 4 or more times weekly | 2.03 | (1.77-2.31) | 2.04 | (1.69-2.46) |
| NHIS: National Health Interview Survey |  |  |  |  |
| VA: Veteran’s Health Administration clinics |  |  |  |  |
| HR: Hazard ratio |  |  |  |  |
| CI: Confidence Interval |  |  |  |  |

**Figure 2:**
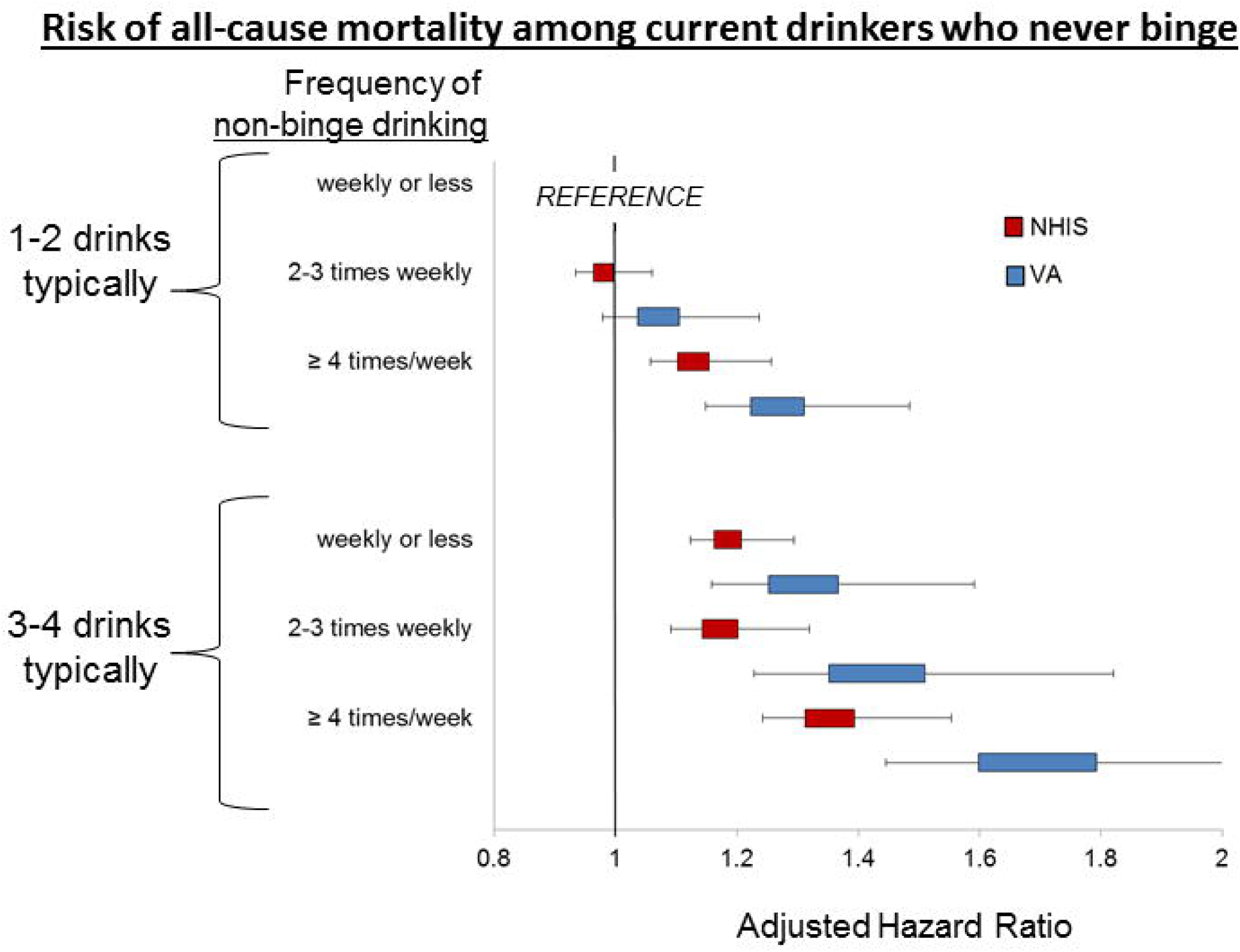
Adjusted hazard ratios for drinking characteristics in non-binge drinkers from survival analysis. Each hazard ratio is adjusted for the other drinking characteristics, age, sex, race/ethnicity, and comorbidity.

In both datasets, the estimated hazard ratios show a statistically significant increase in all-cause mortality for drinking 3-4 drinks when not binging relative to 1-2 drinks when not binging, and for drinking 4 or more times weekly relative to drinking weekly or less (Table 4). Increased mortality is seen for binge drinking more than monthly in the NHIS data and for any amount of binge drinking in the VA data.

A strong potential confounder is smoking because those who drink alcohol are more likely to smoke. Therefore, if smoking is not adequately adjusted for, any increased mortality seen from drinking may be due to smoking. In order to evaluate this, we looked at the association between drinking behavior and mortality only among never smokers in the NHIS data (N=115,325). Similar trends were seen in this subset relative to the full analyses: drinking 4 or more times weekly had increased mortality relative to drinking less frequently (HR 1.07, 95% CI 0.88-1.30), drinking 3-4 drinks when not binging had increased mortality relative to drinking 1-2 drinks at a time (HR 1.22, 95% CI 1.05-1.41), and, relative to never or rarely binging (monthly or less), binging twice monthly to 3 times weekly had increased risk (HR 1.52, 95% CI 1.19-1.93) and binging 4 or more times weekly had increased risk (HR 2.35, 95% CI 1.58-3.49). This suggests that the observed effects of alcohol use on mortality are not due to smoking.

## Discussion

Consistent with previous studies, the results show an increased risk in mortality for drinking 3-4 drinks on a typical occasion relative to drinking 1-2 drinks on a typical occasion. This is consistent with current U.S. dietary guidelines (U.S. Department of Health and Human Services and U.S. Department of Agriculture, 2015), which suggest limiting alcohol consumption to 2 drinks on an occasion for men and one drink for women.

In addition, contrary to previous findings for cardiovascular health (Mukamal et al., 2003; Mukamal et al., 2005; Tolstrup et al., 2006), which found that drinking a greater number of days per week was associated with decreased risk, our analyses show that there is an increased risk for all-cause mortality among people who drink four or more times weekly, relative to people who drink less frequently. This is true even among individuals who drink 1-2 drinks on occasion and never binge. These findings are in contrast to the current US dietary guidelines that do not specify a limit to drinking frequency. Although previous studies have examined drinking quantity and frequency separately (Britton and Marmot, 2004, Mukamal et al., 2003, Mukamal et al., 2005, Mukamal et al., 2010, Tolstrup et al., 2006), much of the focus of research in this field has been on the average number of drinks per day (Chomistek et al., 2015, Costanzo et al., 2010a, Joosten et al., 2011, Mostofsky et al., 2016, Stockwell and Room, 2012) and/or binge drinking (Plunk et al., 2014, Naimi et al., 2013, Pletcher et al., 2005, 2007), both of which have been found to be associated with all-cause mortality. These results highlight a third dimension, drinking frequency, as independently associated with mortality.

Our analytic strategy sought to minimize the risk that the observed effects of alcohol on mortality are due to confounders. All analyses were adjusted for age, sex, race, and comorbidity. In addition, we adjusted for smoking status in the NHIS data (smoking status was not available in the VA data). Finally, we repeated the analyses in the subset of the NHIS data that reported never smoking. Because the findings were robust across analyses, we concluded that the observed associations are unlikely to be due to these potential confounders. However, due to limitations of the available data, there are other potential confounders, including education, marital status, physical activity, and diet, that were not controlled for.

The datasets used in this study are large and have complementary designs, but there are factors that limit the generalizability. First, both studies relied on in-person measurements of self-reported alcohol use, rather than anonymous reports. Report of alcohol use from in-person surveys relative to anonymous surveys may cause bias towards under-reporting drinking (Del Boca and Darkes, 2003, Polich, 1982). Second, the surveys are from a single time point. This would likely increase variance in measurement, which would decrease power. Finally, this study does not distinguish between all-cause mortality and cardiac mortality. This is a particularly important distinction because there is a large body of literature that supports decreased cardiac mortality among light to moderate drinkers (Ronksley et al., 2011, Mukamal, 2007, Gepner et al., 2016, Brien et al., 2011, Mukamal et al., 2010).

To our knowledge, this is the first report of an association between increased mortality and drinking behaviors that fall within the current guidelines for “healthy” alcohol use. Consuming 1-2 drinks at a time on four or more occasions weekly was associated with elevated risk of all-cause mortality, relative to drinking less frequently. This finding was observed in both a large epidemiological sample and a large clinic population.

## Acknowledgements

This research was funded by grants from the National Institutes of Health: K08 DA032680 and R21 AA024888 to SMH, R01 DA036583 to LJB, and UL1 RR024992 to BE. This research is the result of work supported with resources and the use of facilities at the VA Eastern Kansas Healthcare System (Leavenworth, Kansas campus).

---

**Supplemenatry Table S1:** Categorization of non-binge frequency in the VA data.

|  |  | Overall drinking frequency (AUDIT-C item 1) |  |  |  |  |
| --- | --- | --- | --- | --- | --- | --- |
|  |  | Never | Monthly or less | 2-4 times monthly | 2-3 times weekly | ≥ 4 times weekly |
| Binge drinking frequency (AUDIT-C item 3) | Never | No moderate drinking | Weekly or less | Weekly or less | 2-3 times weekly | ≥ 4 times weekly |
|  | < monthly |  | No moderate drinking | Weekly or less | 2-3 times weekly | ≥ 4 times weekly |
|  | Monthly |  | No moderate drinking | Weekly or less | 2-3 times weekly | ≥ 4 times weekly |
|  | Weekly |  |  | No moderate drinking | 2-3 times weekly | ≥ 4 times weekly |
|  | Daily/almost daily |  |  |  | No moderate drinking | No moderate drinking |

**Supplemenatry Figure S1:**
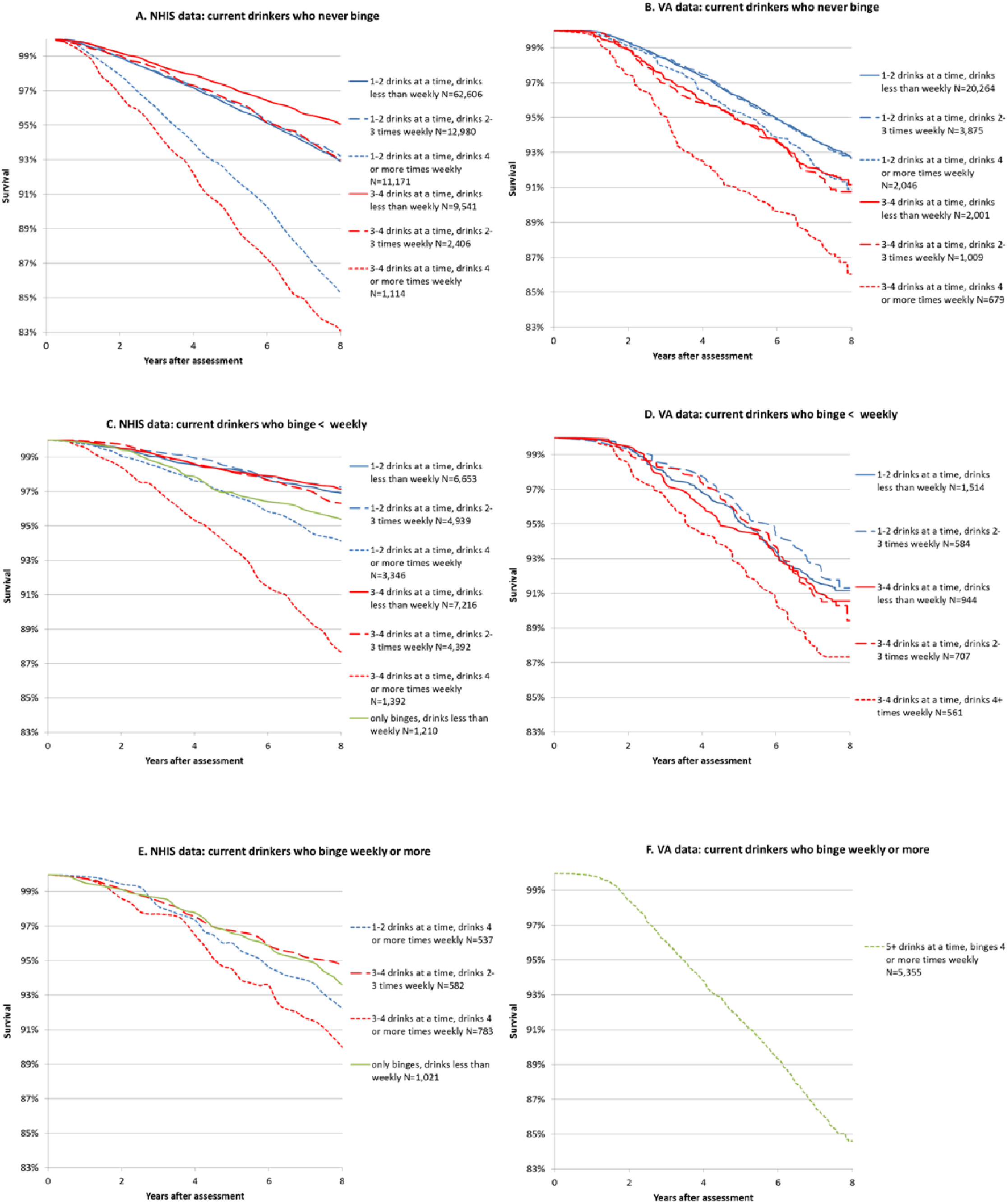
Survival curves stratified by binge drinking behavior. Subgroups with less than 500 people are not included in the plots.

**Supplementary Figure S2:**
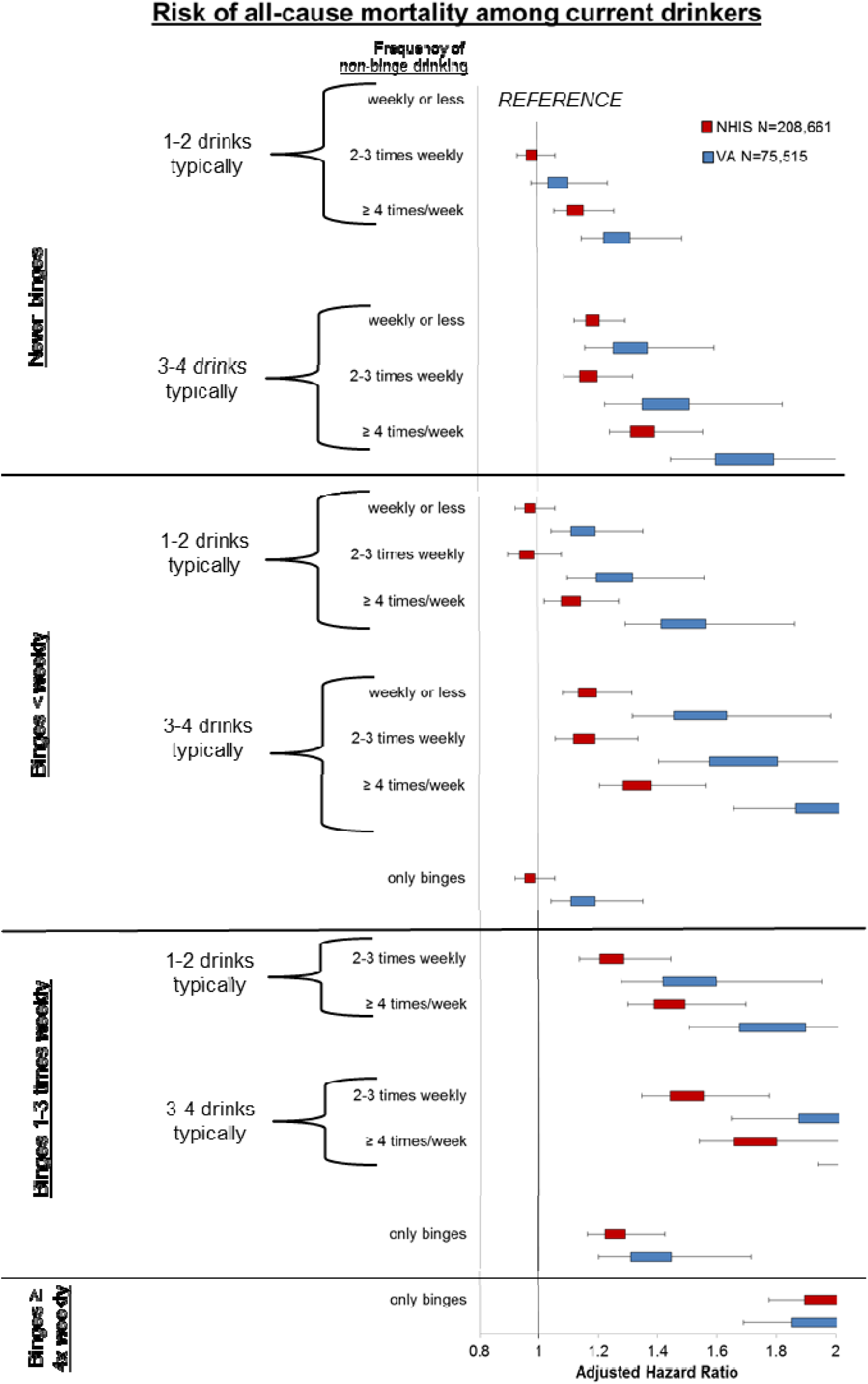
Adjusted hazard ratios for drinking characteristics from survival analysis. Each hazard ratio is adjusted for the other drinking characteristics, age, sex, race/ethnicity, and comorbidity.

